# Metaproteomic responses of *in vitro* gut microbiomes to resistant starches: the role of resistant starch type and inter-individual variations

**DOI:** 10.1101/2020.02.28.970186

**Authors:** Leyuan Li, James Ryan, Zhibin Ning, Xu Zhang, Janice Mayne, Mathieu Lavallée-Adam, Alain Stintzi, Daniel Figeys

## Abstract

Resistant starches (RS) are dietary compounds processed by the gut microbiota into metabolites, such as butyrate, that are beneficial to the host. The production of butyrate by the microbiome appears to be affected by the plant source and type of RS as well as the individual’s microbiota. In this study, we used *in vitro* culture and metaproteomic methods to explore the consistency and variations in individual microbiome’s functional responses to three types of RS - RS2(Hi Maize 260), RS3(Novelose 330) and RS4(Fibersym RW). Results showed that RS2 and RS3 significantly altered the levels of protein expression in the individual gut microbiomes, while RS4 did not result in significant protein changes. Significantly elevated protein groups were enriched in carbohydrate metabolism and transport functions of families Eubacteriaceae, Lachnospiraceae and Ruminococcaceae. In addition, Bifidobacteriaceae was significantly increased in response to RS3. We also observed taxon-specific enrichments of starch metabolism and pentose phosphate pathways corresponding to this family. Functions related to starch utilization, ABC transporters and pyruvate metabolism pathways were consistently increased in the individual microbiomes in response to RS2 and RS3; in contrast, the downstream butyrate producing pathway response varied. Our study confirm that different types of RS have markedly variable functional effects on the human gut microbiome, and also found considerable inter-individual differences in microbiome pathway responses.

## Introduction

Prebiotics are functional compounds that modulate the gut microbiome, promoting the growth and activity of bacteria that are beneficial to human health[1]. This can enhance immune system function and protect from diseases[2]. Resistant starches (RS) are prebiotic polysaccharides that resist digestion by pancreatic amylase and are therefore not hydrolyzed to D-glucose in the small intestine. Current perspectives differ on the definitions and number of RS structural types[3–7]. Resistant starch occurs naturally in three different forms (RS1/RS2/RS3), and synthetically as fourth (RS4)[4–7] and fifth (RS5) form[3, 5]. Because they resist digestion in the small intestine, RS can reach the colon, where they can be fermented by the gut microbiome[8, 9]. RS have been linked to a number of host-beneficial effects when included in human diets[10].

RS can alter the taxonomic composition and functions of the gut microbiome; however, mixed and sometimes opposite effects have been reported [11–15]. In one study, an increase of the Bacteroidetes phylum relative to Firmicutes phylum was observed following an RS4-enriched diet and the opposite following an RS2-enriched diet in human individuals with potential metabolic syndrome [15], whereas in another study RS2 and RS4 in the diet resulted in vastly different effects on the composition of the gut microbiota in healthy human subjects[16]. Nevertheless, both RS types induced physiological changes that were highly similar[16]. In bacterial monoculture, RS3 from two different plants had different effects on short-chain fatty acid (SCFA) production levels in *Clostridium butyricum* and *Eubacterium rectale*[17]. Moreover, it appears that microbiome variations among individuals significantly alter the response to RS. For example, fecal butyrate levels varied widely among 46 individuals given an RS dietary supplement[18]. In another study, MSPrebiotic (containing 70% RS2) increased SCFA levels in elderly adults but not mid-age adults, despite a significant increase in *Bifidobacterium* in both age groups [11]. Overall, the effects of RS on the microbiome appears to depend on the source and type of RS, and on individual variations in the human gut microbiomes. Recent studies on microbiome responses to RS are primarily based on metagenomics and metabolomics approaches[9, 19, 20]. The functional response of the microbiome to different types of RS has remains largely unexplored at the protein level.

We previously developed an *in vitro* culturing method to facilitate individualized evaluation of stimulus effects on the gut microbiome[21]. In our current study, we used this culturing method to evaluate the effect of three types of RS, namely RS2 (enzymatically-resistant starch), RS3 (retrograded starch) and RS4 (chemically-modified starch), on the functionality of *in vitro* gut microbiomes from seven healthy individuals. Through metaproteomics, protein-level variations between individual microbiomes responding to RS types were elucidated in this study. These variations suggest that the functional profiles of microbiomes differ significantly in the presence of RS, and these differences depend on two factors: the RS treatment that a microbiome is responding to, and the composition of an individual’s specific microbiome.

## Results

### Overview of microbiome response to resistant starches

We explored whether functional changes in individual microbiomes induced by RS are dependent on the type of starches and on individual microbiomes. Briefly, stool microbiomes from seven healthy individuals were cultured for 24 hr in the presence of RS or controls using our previously described method[21] (Figure 1). The microbiomes were treated with RS2(Hi Maize 260), RS3(Novelose 330), RS4(Fibersym RW), a non-resistant starch control (corn starch (CS)), a positive control (fructo-oligosaccharide (FOS) known to consistently and markedly shift the *in vitro* gut microbiome[22–24]) or a blank control containing no compounds. The samples were then subjected to metaproteomic analysis as previously described[21]. Altogether, the bioinformatic analysis of 157 LC-MS/MS raw files by MetaLab[25] identified 5,119,332 MS/MS spectra, 80,297 peptides and 21,240 protein groups using a false discovery rate (FDR) threshold of <1% (Supplementary Figure S1). There were 20,378 protein groups with at least one type of functional annotation; 93% had clusters of orthologous groups (COG) annotation, and 70% had Kyoto encyclopedia of genes and genomes (KEGG) annotation. MetaLab identified 142 bacterial species with ≥ 3 peptide assignments (corresponding to 79 genera and 16 phyla).

**Figure 1.**
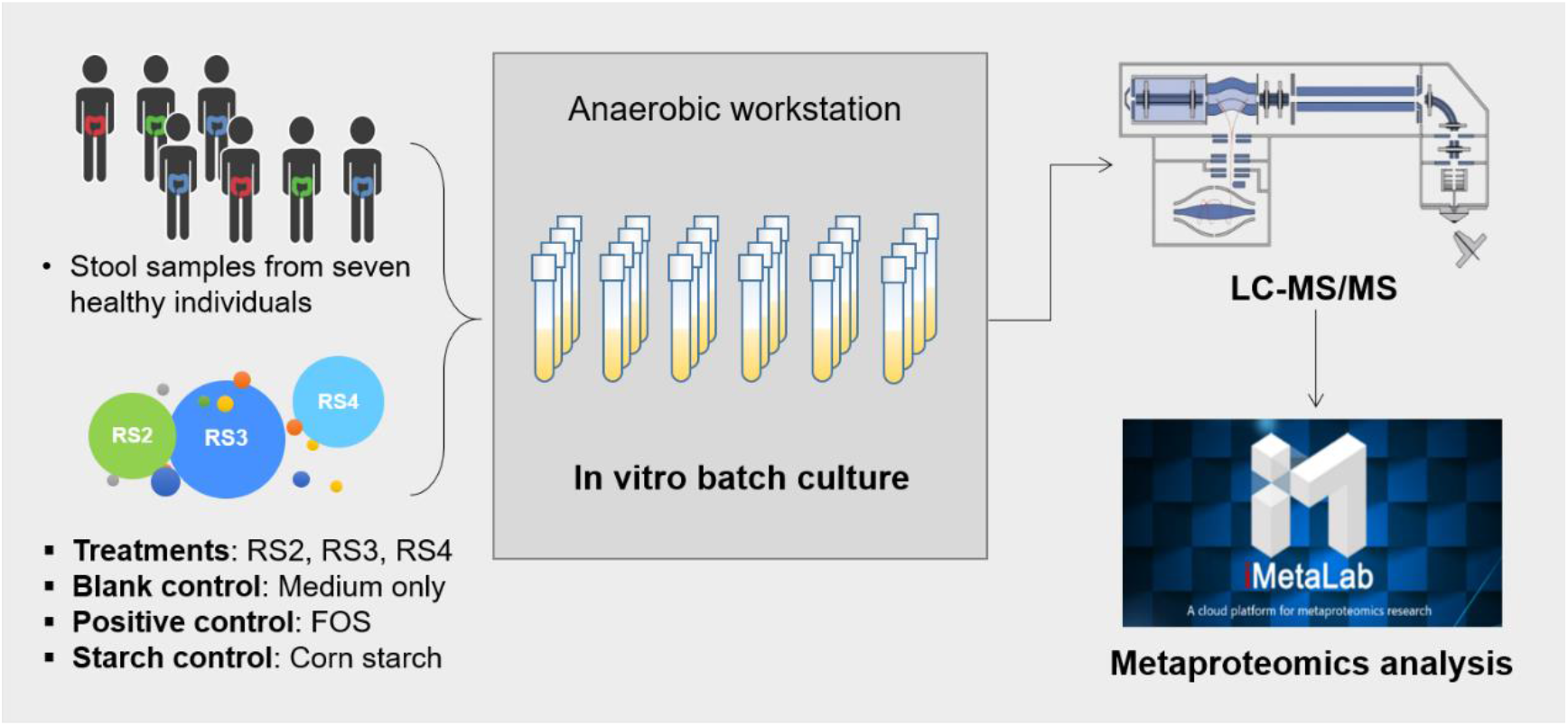
Experimental design. Fecal microbiomes from seven individuals were cultured in medium (as the blank control) or culture medium containing one of the following materials: RS2 (Hi Maize 260), RS3(Novelose 330), RS4(Fibersym RW), FOS, or corn starch. Cultured microbiomes were analyzed using our metaproteomics workflow.

Principle component analysis (PCA) revealed that each individual’s RS treated samples clustered closely with the blank control and CS treatment, whereas FOS treatment, as expected, consistently shifted the microbiome metaproteomic profile for all individual microbiomes (Figures 2A and 2B). In our dataset, the protein groups were annotated to 24 COG categories (Figure 2D). The treatment with the positive control FOS showed 16 COG categories significant changed (total abundance of their protein constituents compared to the blank control). This was consistent with our previous experiments with FOS[24]. The corn starch-treated microbiomes only had three significantly changed COG categories whereas microbiomes treated with the three RS showed a total of 12 significantly changed COG categories of which three COG categories were only changed in the presence of RS.

**Figure 2.**
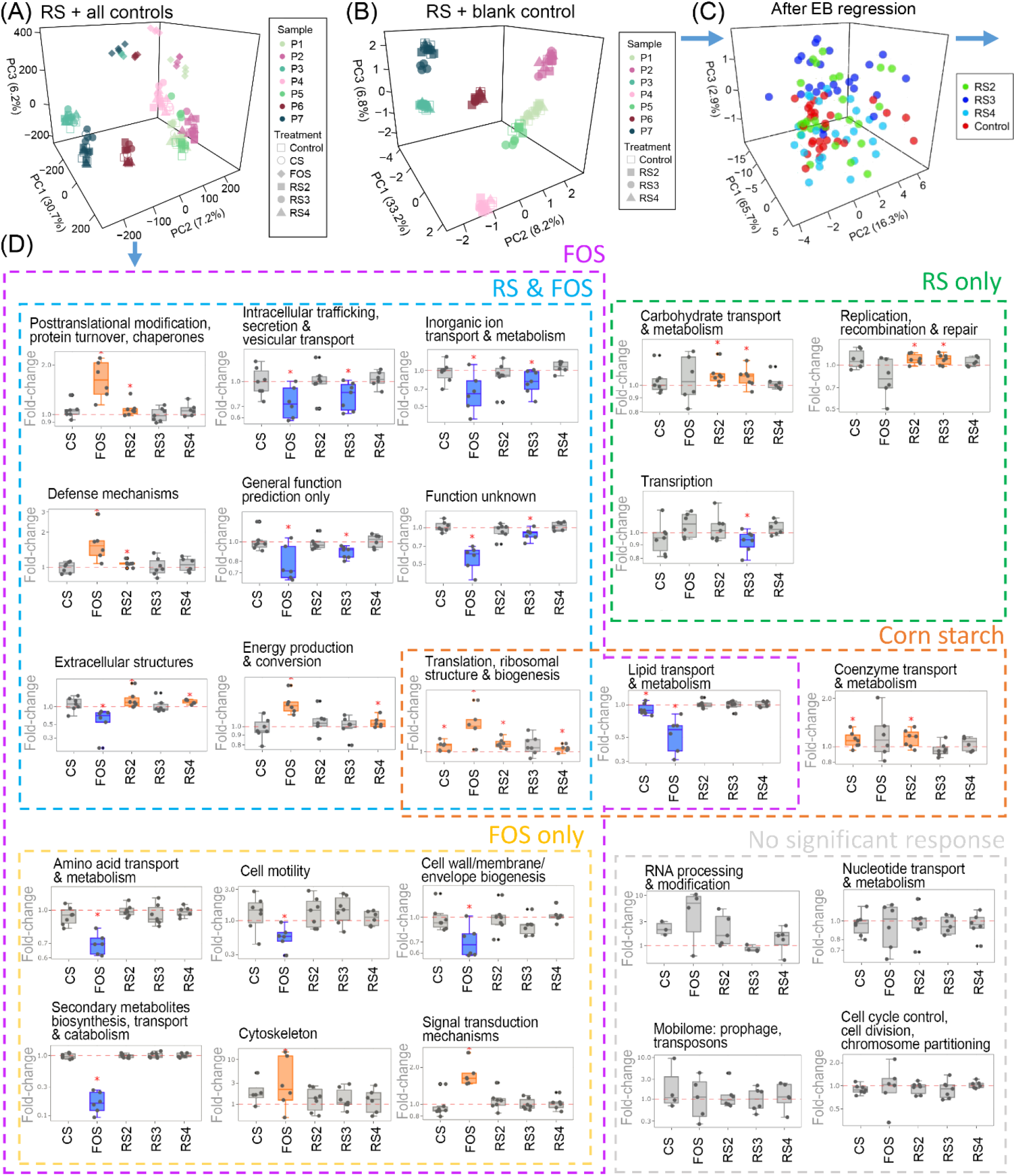
Overview of microbiome response to RS and controls. (A) PCA generated using RS-treated, FOS-treated, CS-treated and blank control microbiomes; (B) PCA generated using RS-treated and blank control microbiomes; (C) PCA of RS-treated and blank control microbiomes after EB regression; (D) Response of COG categories to RS, FOS and CS, in comparison to the blank control sample; the scale of y-axis was log10-transformed. Significant responses (Wilcoxon test, *p* <0.05) were marked with “*”, significant increases are plotted in orange boxes and significant decreases are plotted in blue boxes. Dashed frames gathered significant changes that occurred under the treatment(s), labeled above each colored frame. Blue arrows indicate logical relationship during the data analysis; blue arrow derived from (C) directs to Figure 3A.

As expected, the largest contributors to the differences observed by PCA (Figure 2A and 2B) were the individual microbiomes. We applied an empirical Bayesian (EB) regression approach[26] to remove the inter-individual variance as well as possible batch effects which overshadowed the responses to RS (Figure 2C). This approach allows for the combination of multiple balanced experiments[26] and is robust to small sample sizes. By separating each individual’s microbiome cultures as a subgroup, the experimental design was balanced between subgroups. After EB correction, the inter-individual variance in our dataset was removed, and all subgroups were centered on the PCA (Figure 2C). A clear separation between RS3 and Control was observed, while RS2 and RS4 didn’t show obvious modulating effects on in vitro microbiome.

### Different resistant starches lead to different metaproteomic responses

We then further examined the effect of each RS on the microbiome metaproteomic profiles based on the EB-processed data. PCA using control and each RS types individually revealed that RS2 and RS3 altered the metaproteomic profiles compared to the blank control (Figure 3A and B), whereas RS4-treated samples did not separate from the control (Figure 3C). It was also clear from hierarchical clustering analysis (Supplementary Figure S2) that samples treated by RS2 and RS3 were distinct from RS4 which clustered with the control.

**Figure 3.**
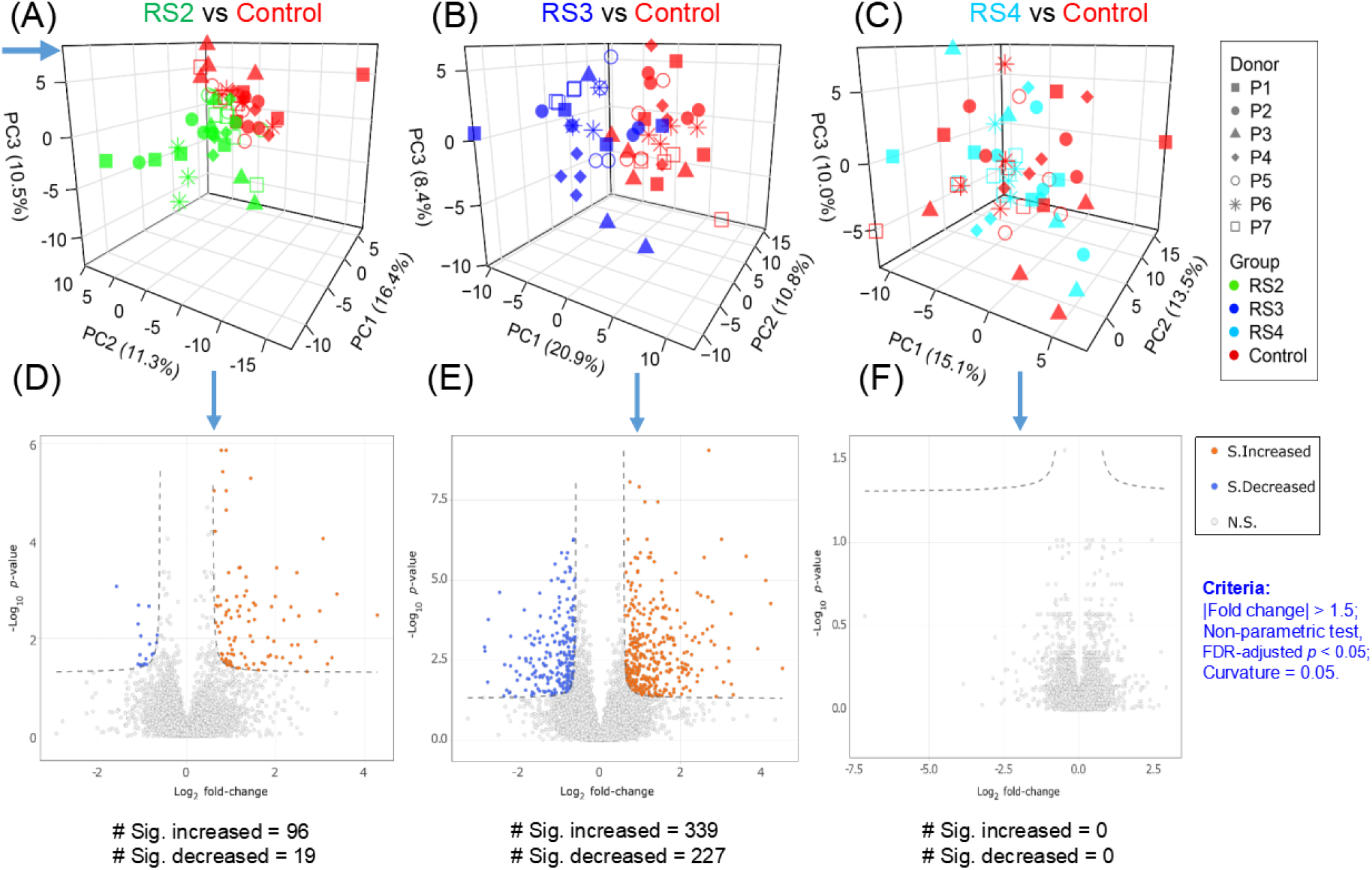
Analysis of differentially expressed protein groups. (A)-(C) PCA plots of RS2-RS4 versus the blank control using EB-corrected data; (D)-(F) Volcano plots of RS2-RS4 versus the blank control generated using EB-corrected data. The *p* values were calculated using Wilcoxon test, and were adjusted using FDR method. Blue arrow means logical relationship in the data analysis workflow; blue arrow pointing to (A) derived from Figure 2C.

Analysis of the differentially expressed protein groups between each RS treatment and the control revealed that RS2 led to 96 protein groups with a significantly increased expression, and 19 with a significantly decreased expression (Figure 3D). In contrast, RS3 resulted in 339 significantly increased and 227 significantly decreased protein groups (Figure 3E). Since RS4 did not show any significant protein expression response (Figure 3F), our subsequent analyses focused on RS2 and RS3.

Taxon-specific functional enrichment (*p* < 0.05) of the proteins significantly increased by RS2 revealed an enrichment in carbohydrate metabolism & transport, amino acid metabolism & transport, as well as translation pathways of butyrate producers that belongs to families Eubacteriaceae and Lachnospiraceae (Figure 4A and Supplementary Figure 3A). Enrichment of carbohydrate metabolism & transport, and translation pathways were also found with RS3. In addition to bacteria from families Eubacteriaceae and Lachnospiraceae, families Bifidobacteriaceae, bacteria from Ruminococcaceae and Bacteroidaceae also contributed to these functional enrichments in response to RS3(Figure 4B and Supplementary Figure 3B).

**Figure 4.**
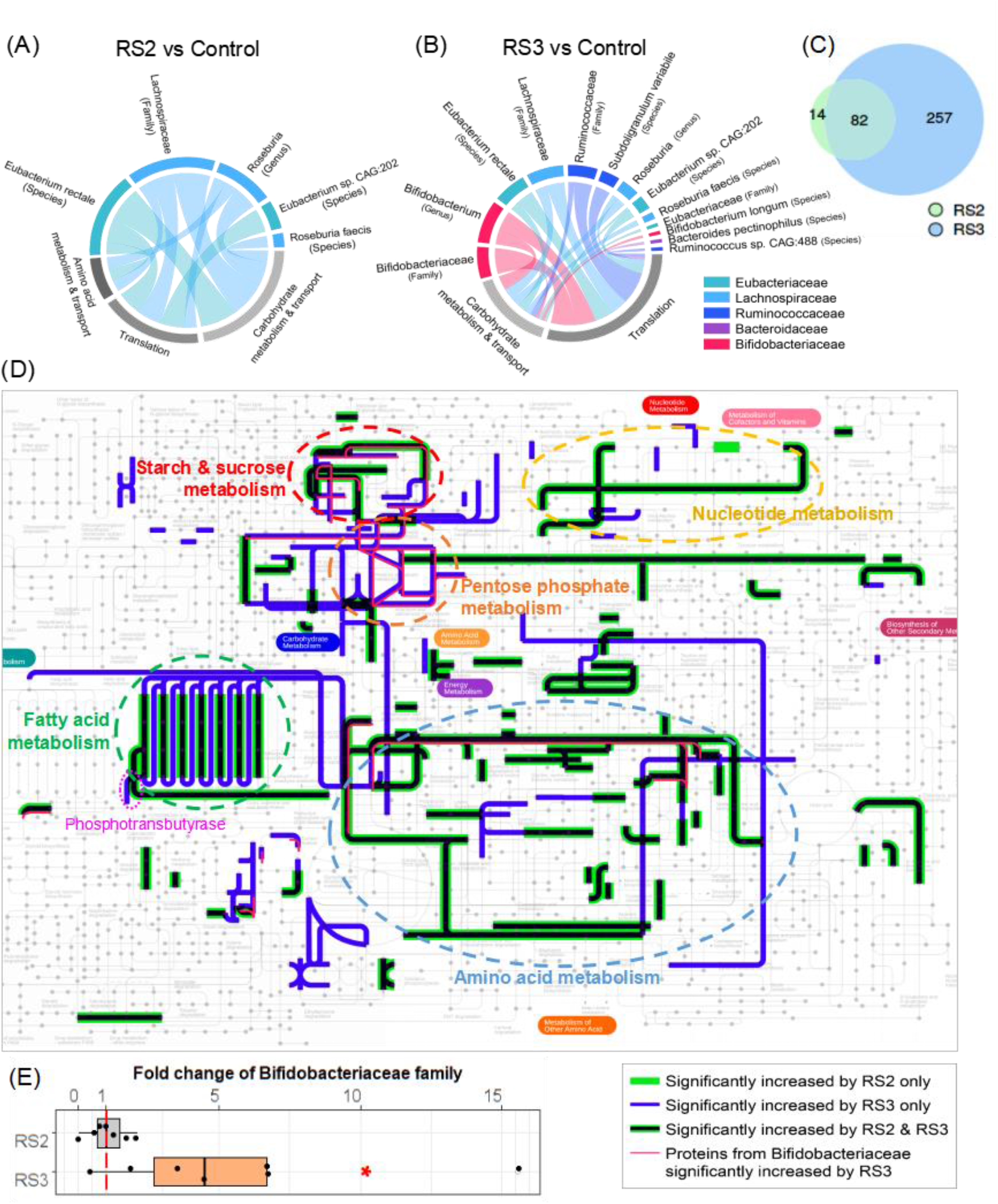
Functional and taxonomic profiles of the significantly increased proteins. (A) Taxon-specific functional enrichment of the significantly increased protein groups in response to RS2; (A) Taxon-specific functional enrichment of the significantly increased protein groups in response to RS3; (C) Comparison of significantly increased protein groups between RS2- and RS3-treated groups; (D) pathways of the significantly increased protein groups; (E) fold change of the Bifidobacteriaceaefamily in RS2- and RS3-treated groups. Red asterisk indicates statistical significance using one-sided Wilcoxon test (*p* < 0.05).

Interestingly, we found that most of the significantly upregulated proteins from RS2 (82 out of 96) were also up-regulated by RS3 (Figure 4C). We mapped COG numbers corresponding to these significantly up-regulated protein groups to iPath 3 (https://pathways.embl.de/)[27]. We merged the result of RS2 and RS3(Figure 4D), the pathway changes caused by RS3 overlapped completely with those cause by RS2 (with only one exception in nucleotide metabolism). Commonly upregulated pathways in both RS groups included starch and sucrose metabolism, nucleotide metabolism, fatty acid metabolism and amino acid metabolism. Notably, proteins involved in pentose phosphate metabolism were only up-regulated by RS3, and more pathways of fatty acid metabolism were upregulated by RS3. Phosphotransbutyrase, a key enzyme in butyrate synthesis[28, 29], was significantly upregulated only by RS3.

The probiotic bacterial family Bifidobacteriaceae (as well as its genus *Bifidobacterium* and species *Bifidobacterium longum*) was only enriched in protein groups upregulated by RS3, and it was the most significantly enriched family in the list of RS3 responders (Supplementary Figure S3). We compared the protein biomass between the control and RS groups based on summed peptide intensities unique to this family. RS3 showed an average fold-change of 5.6 compared to the control (Figure 4E), which was significant (*p* < 0.05) by one-sided Wilcoxon test. We overlaid the protein groups corresponding only to Bifidobacteriaceae to iPath 3 (Figure 4D, red line). This showed that Bifidobacteriaceae mainly contributed in starch and sucrose metabolism and pentose phosphate metabolism pathways, as well as part of amino acid metabolism pathway.

### Inter-individual variation in metabolic pathway responses

We next explored whether the functional changes in response to RS treatments are common across all individual microbiomes or specific to subgroups of microbiomes. We first looked at starch and sucrose metabolism, ABC transporters, and pyruvate metabolism pathways which were significantly increased by RS2 and RS3 treatment (Figure 5A-C). Both RS increased all three pathways in most of the microbiomes: 6/7 microbiomes showed increased starch sucrose metabolism (Figure 5D), 7/7 microbiomes showed increased ABC transporters (Figure 5E); and 6/7 microbiomes showed increased pyruvate metabolism pathways (Figure 5F). There were consistent responses of each downstream metabolic pathway between technical replicates, supporting the reproducibility of the experiment.

**Figure 5.**
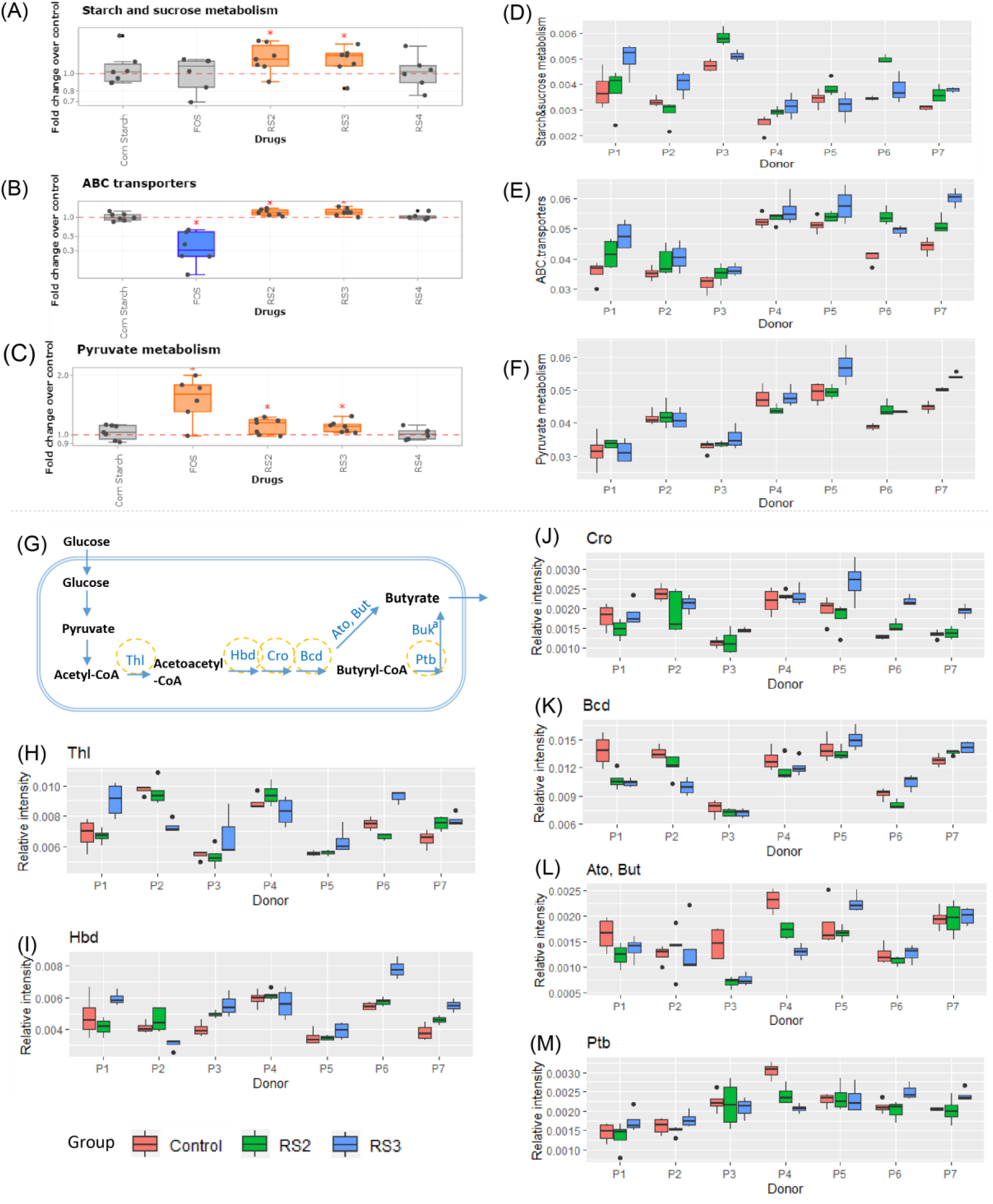
Metabolic pathway responses at an individual resolution. (A)-(C) fold change of starch metabolism, ABC transporters and pyruvate metabolism in all samples. (D)-(F) individual microbiome-specific view of starch metabolism, ABC transporters and pyruvate metabolism responses to RS2 and RS3. (G) Acetyl-CoA pathway for butyrate synthesis (Thl, thiolase; Hbd, β-hydroxybutyryl-CoA dehydrogenase; Cro, crotonase; Bcd, butyryl-CoA dehydrogenase; Ato, butyryl-CoA:acetoacetate CoA transferase (α, β subunits); But, butyryl-CoA:acetate CoA transferase; Ptb, phosphate butyryltransferase; Buk, butyrate kinase); (H)-(M) individual microbiome responses of enzymes involved in butyrate synthesis. (P1-P7 represent seven individual microbiomes, box plots were generated using n=4 technical replicates). ^a^ Buk was not consistently measured across all microbiomes.

Finally, we investigated whether the downstream butyrate production pathway had similar profiles across the individual microbiomes. Interestingly, enzymes within the acetyl-CoA pathway of butyrate production showed different responses(Figure 5G)[30]. RS3 increased Thl (thiolase), Hbd (β-hydroxybutyryl-CoA dehydrogenase) and Cro (crotonase) in 5/7 individual microbiomes, whereas RS2 showed no inter-individual consistency (Figure 5H-J). Of the remaining 2 microbiomes, P2 showed decreases, and P4 did not show apparent responses for these enzymes (Figure 5H-J). In contrast, the downstream enzymes Bcd (butyryl-CoA dehydrogenase), Ato (butyryl-CoA:acetoacetate CoA transferase (α, β subunits)) & But(butyryl-CoA:acetate CoA transferase) and Ptb (phosphate butyryltransferase) did not show inter-individual consistency (Figure 5K-M). Five out of seven individuals increased in overall Ptb, but the changes in Ptb were not statistically significance at the cohort level. Finally, two microbiomes, P6 and P7), showed increases in all the enzymes along the pathway whereas one microbiome (P2) showed primarily decreased levels of enzymes following RS treatments. Butyrate kinase was not consistently measured across all microbiomes and was therefore excluded in the analysis.

## Discussion

In this study, we investigated the functional and taxonomic changes in individual microbiomes exposed to different type and sources of RS using *in vitro* culturing and metaproteomics. The effects of RS on the microbiomes were milder than the changes observed with FOS, which is known to induce taxonomic and functional changes in the human gut microbiome *in vitro*[22–24]. The microbiomes treated with RS maintained their individuality (Figure 2A and B) whereas FOS treatment tended to regroup the microbiomes (Figure 2A). Although the PCA showed only weak changes induced by RS treatments, we still observed many significantly shifted microbiome function in response to RS (RS2 and RS3), both by using paired comparisons for individual microbiomes (Figure 2D), and by using EB-transformed protein intensities (Figure 3). Interestingly, RS2 and RS3 are both derived from high amylose maize and had similar functional effects of increasing carbohydrate metabolism and transformation in butyrate producing bacteria from families Eubacteriaceae and Lachnospiraceae (Figure 4A and B). Nevertheless, RS3 had higher number of enriched taxa corresponding to its functional response, and significantly increased proteins were found in Bifidobacteriaceae (Supplementary Figure S3). Proteins from Ruminococcaceae and Bacteriodiaceae were also increased in response to RS3. In agreement with our findings, previous studies have reported increases of bacteria from families Bifidobacteriaceae [11, 12, 31–33], Ruminococcaceae[12, 13, 15, 33], Eubacteriaceae[15, 33, 34], Lachnospiraceae[35] and Bacteriodiaceae[15, 32] in response to different sources and forms of RS. Members of these bacterial families have been shown to be capable of metabolizing resistant starches[17, 36–38] due to their amylolytic activities[36, 38], and many are butyrate producers[39]. Our findings showed that the protein groups up-regulated by RS2 and RS3 were enriched in starch metabolism pathways. In response to RS3, Bifidobacteriaceae showed increases in the starch uptake and pentose phosphate pathways, which contributed significantly to the overall increase seen in the pentose phosphate pathway. The product of the pentose phosphate pathway, NADPH, is a major source of electrons for diverse anabolic biosynthetic processes, and therefore a primary reducing agent for biosynthesis pathways[40], including fatty acid metabolism and especially butyrate production. However, our results showed that the response of the pentose phosphate pathway was strongly related to the type of the RS, indicating that different types of RS may result in various effects on biosynthesis pathways including butyrate generation. Although RS2 and RS3 were both derived from high-amylose starch, they are not chemically identical (RS2 – hydrothermal treated, RS3 - retrogradation), which may contribute to the difference of functional and taxonomic responses observed in our study. RS4 is a type of chemically modified starch that is not naturally occurring. In this study, we did not observe significant protein changes induced by RS4, which is similar to a recent study reporting that an RS4 did not affect the microbiome[41]. In agreement with a single strain-based study on *Eubacterium rectale*[38], we found that microbiome functions of starch utilization and ABC transporters were increased in the presence of RS. The gut bacteria expressed enzymes for starch degradation, and the ABC glycan-binding proteins were increased to scavenge the liberated maltooligosaccharides and glucose[38]. Subsequently, the pyruvate metabolism pathways utilizing glucose were also significantly increased. These pathways were consistently increased across all individual microbiomes. Interestingly, derived from pyruvate, the downstream acetyl-CoA pathway (Figure 5G) of butyrate production started to show inter-individual differences. Responses of the acetyl-CoA pathway to RS2 were largely inconsistent among different individuals’ microbiomes. RS3, which showed stronger effects than RS2, consistently increased enzymes along the acetyl-CoA pathway in specific individual microbiomes, while other microbiomes had decreases along this pathway (Figure 5H-M). McOrist *et al* observed that fecal butyrate levels vary widely among individuals when supplemented with an RS diet [18]. In agreement with their findings, our results provided a pathway view of how the butyrate production enzymes respond differently in individual microbiomes. Although our data showed that protein groups involved in starch metabolism and related pathways were initially significantly increased in butyrate producers, the butyrate producing pathway response varied widely. This may be due to different forms of inter-specific interactions within individuals’ microbial ecosystems or differences in response time across microbiomes, and is potentially indicative that some microbiomes were not increasing their production of butyrate following exposure to specific RS.

In summary, our study demonstrated that different types of RS have markedly variable functional effects on the human gut microbiome. Even if the source of RS is the same, different ways of forming the RS can result in significantly different responses. We also demonstrated that inter-individual differences in microbiome pathway responses were considerable. There are numerous types of marketed RS products, which are derived from various plant sources and are processed in to RS in different ways. Our study suggested the importance of individualized screening of dietary RS, using high-throughput approaches like RapidAIM assay [42], for improving health or for their utilization as personalized intervention for human diseases.

## Materials and methods

### Resistant starches and controls

RS involved in this study were, RS2 [Hi Maize 260, Ingredion, Inc., Westchester, IL, USA], containing 60% of RS2; RS3 [Novelose 330, obtained from Ingredion, Inc., Westchester, IL, USA], containing 28-38% of RS3; and RS4 [Fibersym RW, obtained from MGP Ingredients, Atchison, KS, USA], containing 85% RS4. Due to the fact that the RS contain different proportion of non-resistant starch, we included corn starch as the negative control. Fructo-oligosaccharide (FOS) - Orafti P95 (BENEO, Inc., Parsippany, NJ, USA), known to consistently and markedly shift the *in vitro* gut microbiome[22, 23], was used as the positive control. Previous literature involving microbiome responses to prebiotics using fecal slurries inoculations with a 1% w/v prebiotic concentration[43, 44]; this concentration was used for this study’s culture samples. 0.04 g (1% w/v) of RS/corn starch/FOS was added to each corresponding culture tube. For a blank control, microbiome samples were cultured in the same medium but without any compounds added.

### Stool sample collection

This study was following a human stool sample collection protocol (#2016-0585-01H) that is authorized through the University of Ottawa’s Ottawa Health Science Network Research Ethics Board. Stool samples from seven healthy volunteers (22 - 39 years of age; males and females) were involved in this study. Exclusion criteria were: IBS, IBD, or diabetes diagnosis; antibiotic use or gastroenteritis episode in the last 3 months; use of pro-/pre-biotic, laxative, or anti-diarrheal drugs in the last month; or pregnancy. Each potential participant was assigned a number to blind researchers from sample identity. On the day of collection, the participant is given a kit containing a 50 mL BD Falcon™ tube containing 12.5 mL of pre-reduced PBS [311-010-CL; Fisher Scientific, Fair Lawn, NJ, USA] (had been placed in an anaerobic chamber 24 hours before collection) and 0.0125 g of L-cysteine [C7352; Sigma-Aldrich Canada Co., Oakville, ON, Canada] (0.1% w/v; added just before sample collection). Approximately 3 g of fresh stool sample was collected by each participant into buffer described above and returned to study coordinator within 30 min. Samples were then weighed and transferred into an anaerobic workstation (5% H2, 5% CO2, and 90% N2 at 37°C), vortex mixed and filtered through sterile gauzes to remove solids.

### *In vitro* culturing and prebiotic treatment

The microbiome inoculums were treated with the RS2/RS3/RS4/FOS samples through *in vitro* culturing using a bioanalytical assay testing, as previously reported culture medium[21]. Each participant’s stool sample was cultured in 21 separate tubes: four technical replicates for each of the five different treatment conditions (RS2, RS3, RS4, FOS, and the blank), and one well for corn starch as a negative control, with an in-solution stool concentration of 2% w/v. In each culture tube, 4 mL of a basic nutrient and salt medium was added prior to stool inoculation. This medium consisted of peptone water [70179; MilliporeSigma, Oakville, ON, Canada] (0.2 g; 0.2% w/v), yeast extract [212750; BD Biosciences, Sparks, MD, USA] (0.2 g; 0.2% w/v), monopotassium phosphate [P5655; MilliporeSigma] (0.045 g; 0.045% w/v), dipotassium phosphate [PX1570-1; EMD Millipore, Etobicoke, ON, Canada] (0.045 g; 0.045% w/v), sodium hydroxide [S8045; MilliporeSigma] (0.09 g; 0.09% w/v), sodium bicarbonate [SX0320-3; EMD Millipore] (0.4 g; 0.4% w/v), magnesium sulphate heptahydrate [230391; MilliporeSigma] (0.009 g; 0.009% w/v), calcium chloride [CX0156-1; EMD Millipore] (0.009 g; 0.009% w/v), bile salts [48305; MilliporeSigma] (0.05 g; 0.05% w/v), Tween 80 [P1754; MilliporeSigma] (200 μL; 0.2% v/v), and distilled water (~90 mL). This 100 mL solution was scaled to account for the total number of samples, autoclaved, corrected to pH ~7, and placed in an anaerobic chamber for 24 hours. Culture tubes were then placed on a shaking platform (300 rpm) inside the anaerobic chamber at 37 ^o^C for 24 hours.

### Cell washing, cell lysis, protein extraction and tryptic digestion

Cultured samples were centrifuged at 300 g/4 °C for 5 min to remove debris. The supernatant was collected and bacterial cells were pelleted by centrifuging for 20 minutes (14,000*g*/4°C). Bacterial pellets were then washed three times using cold PBS (centrifuged at 14,000*g*/4°C). All samples were stored overnight in −80°C, followed by a cell lysis procedure: 200 μL of a lysis buffer (10 mL recipe: urea [U5128; MilliporeSigma] (3.8 g, 38% w/v); 20% SDS [L3771; MilliporeSigma] (2 mL, 20% v/v); 1M Tris-HCl [C4706; MilliporeSigma] (0.5 mL, 5% v/v); ddH_2_O (4 mL, 40% v/v); PhosSTOP^™^ tablet [4906837001; MilliporeSigma] (one tablet); cOmplete mini^™^ [4693124001;; MilliporeSigma] (one tablet)) was added to each sample culture tube. Each sample was then ultrasonicated (25% amplitude) for 30 seconds, placed on ice for 30 seconds, then ultrasonicated again (same setting) for 30 seconds. Sample tubes were then centrifuged for 10 minutes (16,000*g*/8°C). Supernatants were transferred to new 2.0 mL Axygen^®^ tubes, then 1 mL of a - 20°C pre-chilled precipitation solution was added (50% v/v acetone [A949-4; Fisher Scientific, Fair Lawn, NJ, USA]; 50% v/v ethanol [1009; Commercial Alcohols, Tiverton, ON, Canada]; 0.1% v/v acetic acid [A38-212; Fisher Scientific]). Proteins in samples were then precipitated overnight at −20°C.

Protein samples were washed for three rounds using cold (−20°C) acetone. Then, samples were re-suspended in 100 μL of 6M urea buffer (pH=8.0), and a DC^™^ assay [Bio-Rad Laboratories, Mississauga, ON, Canada] was performed to determine protein content. 50 μg of protein from each sample was used for in-solution digestion. 2 μL of dithiothriotol (DTT) [43815; MilliporeSigma] was added to each tube, then tubes incubated with shaking for 30 minutes on an Eppendorf Thermomixer C (800 rpm/56°C). 2 μL of iodoacetamide (IAA) [I1149; MilliporeSigma] was then added to each sample, and incubated in darkness for 40 minutes at rt. 1 μg trypsin [T1426; MilliporeSigma] was then added to digest the proteins with incubation at 37°C for 24 hours. Samples were then acidified to pH ~2-3 using 5% (v/v) formic acid and were desalted using C18 beads [ReproSil120 C18-AQ; Dr. Maisch GmbH, Ammerbuch-Entringen, DE].

### LC-MS/MS analysis

All samples were run on Eksigent 425 nanoHPLC connected to Orbitrap Elite™ Hybrid Ion Trap Mass Spectrometer [Thermo-Fisher Scientific, Waltham, MA, USA] with a 240 min gradient. Peptides were separated with an in-house made column (75 μm i.d. × 15 cm) packed with reverse phase beads [1.9 μm/120 Å ReproSil-Pur C18 resin, Dr. Maisch GmbH, Ammerbuch, Germany]. The samples were loaded at 5% buffer A (0.1% formic acid in H_2_O) and analyzed by a gradient from 5 to 30% (v/v) buffer B (0.1% formic acid, 80% acetonitrile in H_2_O) at a flow rate of 300 nL/min. MS analysis was done with a full MS scan from 350 to 1750 m/z in Orbitrap, followed by data-dependent MS/MS scan of the 20 most intense ions in ion trap. Microbiome samples corresponding to each individual were run on LC-MS/MS in a randomized order. Spectral data were collected as *.RAW files.

### Metaproteomic data processing and statistical analysis

Database search was performed automatically, following the MetaPro-IQ workflow using the MetaLab software (version 1.0)[25], with MaxQuant version 1.5.3.30 involved in the workflow. Carbamidomethyl (C) was set as fixed modifications, and Acetyl (Protein N-term) and Oxidation (M) modifications were included as variable modification in the database search space. The MetaLab database search results allows for protein identification and quantification and serves as a basis for identifying biochemical pathways that are differentially expressed. The output provides massive information on the dataset, including summary, peptides, protein groups, functional and taxonomic data tables, etc.

Data quality check was performed by submitting the summary.txt file to our MaxQuant Quick Summary app (https://shiny.imetalab.ca/MQ_summary/). LFQ intensities of protein groups were filtered with the criteria of having non-zero values in ≥4 samples in each individual subgroup, at least in four of the individual microbiomes. PCA of the protein groups data was then performed using R function prcomp() and visualized using R package “scatterplot3d”. To overcome inter-individual variance that were overriding RS responses, an Empirical Bayesian-based approach was performed with our online tool (https://shiny.imetalab.ca/batcheffect_explorer/). The corrected data was then scaled by sum sample wise, and then differentially expressed protein groups between two conditions were performed using our online iMetaShiny app - Differential Protein Analyzer (https://shiny.imetalab.ca/Volcano_plot/). Criteria and parameters used for statistical test and visualization were: |Fold change| > 1.5, non-parametric test (Wilcoxon test), FDR-adjusted *p* < 0.05, and curvature = 0.05. Here, we applied a smooth curve cut-off, which was first used in proteomics by Keilhauer *et al*[35]. The smooth curve is defined by the following equation: y = curvature / |x-”Log2FoldChangeCutOff”| + “-Log10pValueCutOff”.

Taxonomic and functional enrichment analysis were performed using our online tool Enrichment Analysis (https://shiny.imetalab.ca/metaproteomics_enrichment/), *p* value significances threshold for the enrichment analyses was set as <0.05. Visualization of pathways was performed using COG accession numbers in iPath 3 (https://pathways.embl.de/)[27]. For functional.csv visualization, we wrapped a R Shiny web page (https://leyuan.shinyapps.io/RS_data/) to compare COG and NOG categories, COG, NOG and KEGG accessions and names, as well as GO comparisons between each treatment and the blank control. Each data point represents the average fold-change of all technical replicates corresponding to one individual microbiome and one treatment. A Wilcoxon test was used to evaluate the statistical significance of the fold change in comparison to the blank control. To identify proteins that are unique to each family, we extracted the protein groups corresponding to this family by mapping its unique peptide IDs (in MetaLab.allPepTaxa.csv) to the protein IDs (in proteinGroups.txt). To summarize the contribution of Bifidobacteriaceae’s proteomic biomass relative to the whole microbiome, as a proxy of the relative total biomass, we summed the peptide intensities corresponding to the Bifidobacteriaceae and calculated its proportion of the total peptide biomass in each sample.

## Acknowledgments

This work was supported by the Government of Canada through Genome Canada and the Ontario Genomics Institute (OGI-156 and OGI-149), the Natural Sciences and Engineering Research Council of Canada (NSERC, grant no. 210034), the Ontario Ministry of Economic Development and Innovation (ORF-DIG-14405 and project 13440), a W. Garfield Weston Foundation Grant to A.S., and an NSERC Discovery Grant to M.L.A; L.L. and J.R. were funded by a stipend from the NSERC CREATE in Technologies for Microbiome Science and Engineering (TECHNOMISE) Program.

## Supplementary material

**Figure S1.**
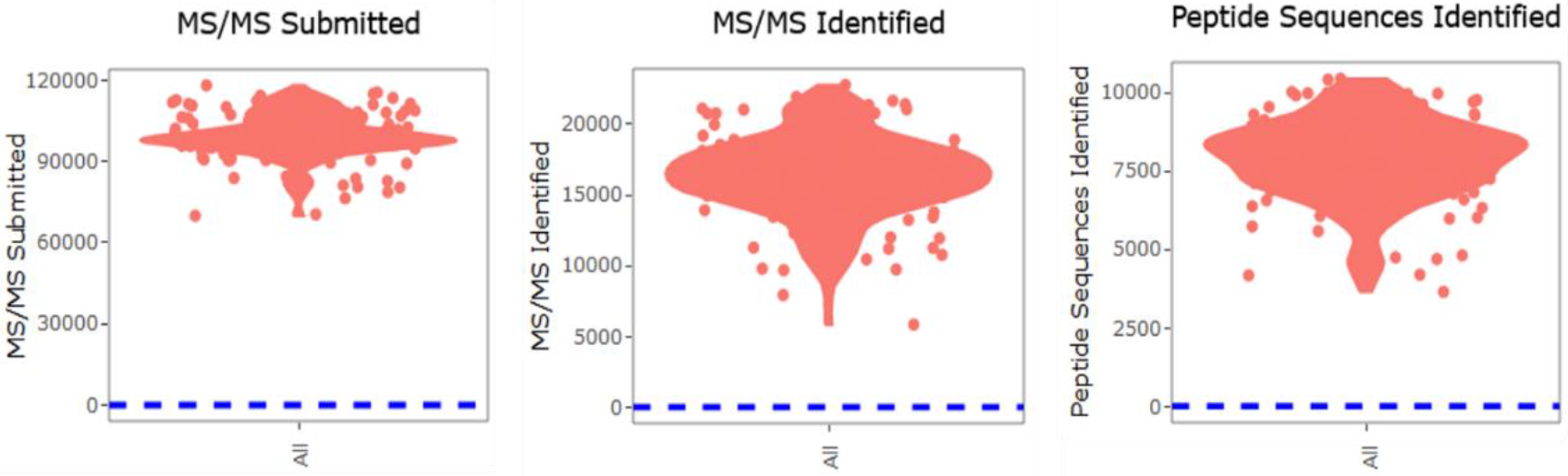
Data quality

**Figure S2.**
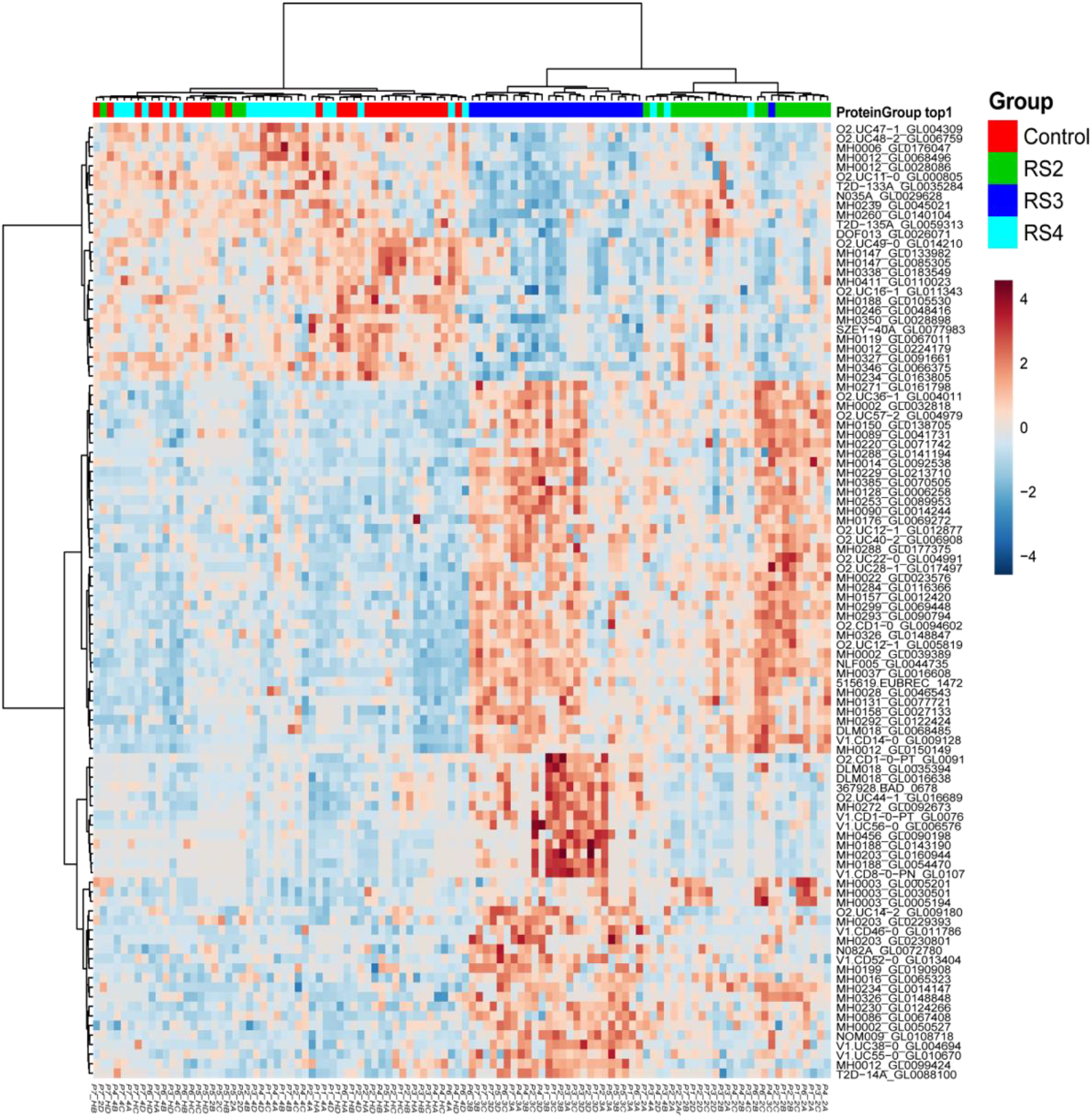
Overall heatmap generated using top 100 ANOVA

**Figure S3.**
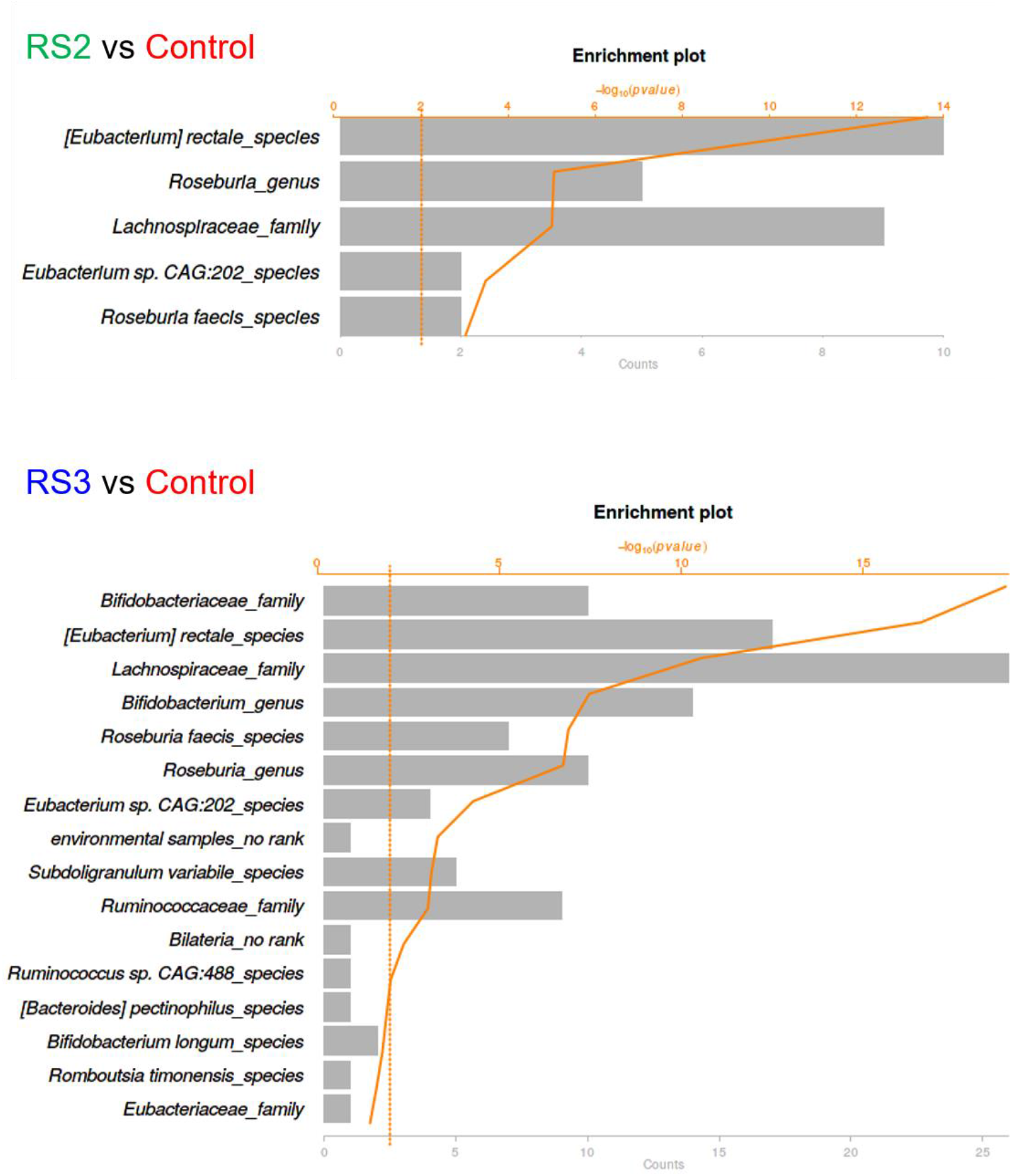
Taxonomic enrichment of the significantly increased proteins

## References

1. Hutkins, R.W., et al., Prebiotics: why definitions matter. Current Opinion in Biotechnology, 2016. 37: p. 1–7.

2. Schley, P.D. and C.J. Field, The immune-enhancing effects of dietary fibres and prebiotics. British Journal of Nutrition, 2002. 87(S2): p. S221–S230.

3. Fuentes-Zaragoza, E., et al., Resistant starch as prebiotic: A review. Starch - Stärke, 2011. 63(7): p. 406–415.

4. Yao, N., A.V. Paez, and P.J. White, Structure and Function of Starch and Resistant Starch from Corn with Different Doses of Mutant Amylose-Extender and Floury-1 Alleles. Journal of Agricultural and Food Chemistry, 2009. 57(5): p. 2040–2048.

5. Birt, D.F., et al., Resistant Starch: Promise for Improving Human Health. Advances in Nutrition, 2013. 4(6): p. 587–601.

6. Sajilata, M.G., R.S. Singhal, and P.R. Kulkarni, Resistant Starch–A Review. Comprehensive Reviews in Food Science and Food Safety, 2006. 5(1): p. 1–17.

7. Higgins, J.A. and I.L. Brown, Resistant starch: a promising dietary agent for the prevention/treatment of inflammatory bowel disease and bowel cancer. Current Opinion in Gastroenterology, 2013. 29(2): p. 190–194.

8. Wang, Y., et al., The Capacity of the Fecal Microbiota From Malawian Infants to Ferment Resistant Starch. Frontiers in Microbiology, 2019. 10.

9. Vital, M., et al., Metagenomic Insights into the Degradation of Resistant Starch by Human Gut Microbiota. Applied and Environmental Microbiology, 2018. 84(23): p. e01562–18.

10. Ashwar, B.A., et al., Preparation, health benefits and applications of resistant starch—a review. Starch - Stärke, 2016. 68(3-4): p. 287–301.

11. Alfa, M.J., et al., A randomized trial to determine the impact of a digestion resistant starch composition on the gut microbiome in older and mid-age adults. Clinical Nutrition, 2018. 37(3): p. 797–807.

12. Kieffer, D.A., et al., Resistant starch alters gut microbiome and metabolomic profiles concurrent with amelioration of chronic kidney disease in rats. American Journal of Physiology-Renal Physiology, 2016. 310(9): p. F857–F871.

13. Maier, T.V., et al., Impact of Dietary Resistant Starch on the Human Gut Microbiome, Metaproteome, and Metabolome. mBio, 2017. 8(5): p. e01343–17.

14. Bergeron, N., et al., Diets high in resistant starch increase plasma levels of trimethylamine-N-oxide, a gut microbiome metabolite associated with CVD risk. British Journal of Nutrition, 2016. 116(12): p. 2020–2029.

15. Upadhyaya, B., et al., Impact of dietary resistant starch type 4 on human gut microbiota and immunometabolic functions. Scientific Reports, 2016. 6(1): p. 28797.

16. Martínez I, K.J., Duffy PR, Schlegel VL, Walter J., Resistant starches types 2 and 4 have differential effects on the composition of the fecal microbiota in human subjects. PLoS One, 2010. 5: p. e15046.

17. Purwani, E.Y., T. Purwadaria, and M.T. Suhartono, Fermentation RS3 derived from sago and rice starch with Clostridium butyricum BCC B2571 or Eubacterium rectale DSM 17629. Anaerobe, 2012. 18(1): p. 55–61.

18. McOrist, A. L., et al., Fecal Butyrate Levels Vary Widely among Individuals but Are Usually Increased by a Diet High in Resistant Starch. The Journal of Nutrition, 2011. 141(5): p. 883–889.

19. Yu, M., et al., Microbiome-Metabolomics Analysis Investigating the Impacts of Dietary Starch Types on the Composition and Metabolism of Colonic Microbiota in Finishing Pigs. Frontiers in Microbiology, 2019. 10(1143).

20. Bang, S.-J., et al., Effect of raw potato starch on the gut microbiome and metabolome in mice. International Journal of Biological Macromolecules, 2019. 133: p. 37–43.

21. Li, L., et al., Evaluating in vitro culture medium of gut microbiome with orthogonal experimental design and a metaproteomics approach. Journal of Proteome Research, 2018. 17(1): p. 154–163.

22. Liu, F., et al., Fructooligosaccharide (FOS) and Galactooligosaccharide (GOS) Increase Bifidobacterium but Reduce Butyrate Producing Bacteria with Adverse Glycemic Metabolism in healthy young population. Scientific Reports, 2017. 7(1): p. 11789.

23. Sivieri, K., et al., Prebiotic Effect of Fructooligosaccharide in the Simulator of the Human Intestinal Microbial Ecosystem (SHIME^®^ Model). Journal of Medicinal Food, 2014. 17(8): p. 894–901.

24. Zhang, X., et al., In Vitro Metabolic Labeling of Intestinal Microbiota for Quantitative Metaproteomics. Analytical Chemistry, 2016. 88(12): p. 6120–6125.

25. Cheng, K., et al., MetaLab: an automated pipeline for metaproteomic data analysis. Microbiome, 2017. 5(1): p. 157.

26. Johnson, W.E., C. Li, and A. Rabinovic, Adjusting batch effects in microarray expression data using empirical Bayes methods. Biostatistics, 2006. 8(1): p. 118–127.

27. Darzi, Y., et al., iPath3.0: interactive pathways explorer v3. Nucleic Acids Research, 2018. 46(W1): p. W510–W513.

28. Walter, K.A., et al., Sequence and arrangement of two genes of the butyrate-synthesis pathway of Clostridium acetobutylicum ATCC 824. Gene, 1993. 134(1): p. 107–111.

29. RC Valentine, R.W., Purification and role of phosphotransbutyrylase. Journal of Biological Chemistry, 1960. 235(7): p. 1948–1952.

30. Vital, M., A.C. Howe, and J.M. Tiedje, Revealing the bacterial butyrate synthesis pathways by analyzing (meta)genomic data. mBio, 2014. 5(2): p. e00889–14.

31. Metzler-Zebeli, B.U., et al., Resistant starch reduces large intestinal pH and promotes fecal lactobacilli and bifidobacteria in pigs. animal, 2019. 13(1): p. 64–73.

32. Gopalsamy, G., et al., Resistant Starch Is Actively Fermented by Infant Faecal Microbiota and Increases Microbial Diversity. Nutrients, 2019. 11(6): p. 1345.

33. Venkataraman, A., et al., Variable responses of human microbiomes to dietary supplementation with resistant starch. Microbiome, 2016. 4(1): p. 33.

34. Verberkmoes, N.C., et al., Shotgun metaproteomics of the human distal gut microbiota. The ISME Journal, 2009. 3(2): p. 179–189.

35. Keilhauer, E.C., M.Y. Hein, and M. Mann, Accurate Protein Complex Retrieval by Affinity Enrichment Mass Spectrometry (AE-MS) Rather than Affinity Purification Mass Spectrometry (AP-MS). Molecular & Cellular Proteomics, 2015. 14(1): p. 120.

36. Ze, X., et al., Ruminococcus bromii is a keystone species for the degradation of resistant starch in the human colon. The ISME Journal, 2012. 6(8): p. 1535–1543.

37. Zhang, Y., et al., The in vitro effects of retrograded starch (resistant starch type 3) from lotus seed starch on the proliferation of Bifidobacterium adolescentis. Food & Function, 2013. 4(11): p. 1609–1616.

38. Cockburn, D.W., et al., Molecular details of a starch utilization pathway in the human gut symbiont Eubacterium rectale. Molecular Microbiology, 2015. 95(2): p. 209–230.

39. Louis, P. and H.J. Flint, Formation of propionate and butyrate by the human colonic microbiota. Environmental Microbiology, 2017. 19(1): p. 29–41.

40. Wolfe, A.J., Glycolysis for Microbiome Generation. Microbiology spectrum, 2015. 3(3): p. 10.1128/microbiolspec. MBP-0014-2014.

41. Deehan, E.C., et al., Precision Microbiome Modulation with Discrete Dietary Fiber Structures Directs Short-Chain Fatty Acid Production. Cell Host & Microbe, 2020.

42. Leyuan Li, Z.N., Xu Zhang, Janice Mayne, Kai Cheng, Alain Stintzi, Daniel Figeys, RapidAIM: A culture- and metaproteomics-based Rapid Assay of Individual Microbiome responses to drugs. BioRxiv:543256, 2019. doi: https://doi.org/10.1101/543256.

43. Rycroft, C.E., et al., A comparative in vitro evaluation of the fermentation properties of prebiotic oligosaccharides. Journal of Applied Microbiology, 2001. 91(5): p. 878–887.

44. Olano-Martin, E., G.R. Gibson, and R.A. Rastall, Comparison of the in vitro bifidogenic properties of pectins and pectic-oligosaccharides. Journal of Applied Microbiology, 2002. 93(3): p. 505–511.

